# Psip1/p52 regulates distal *Hoxa* genes through activation of lncRNA Hottip

**DOI:** 10.1101/061192

**Authors:** Madapura M Pradeepa, Gillian Taylor, Graeme R Grimes, Andrew J Wood, Wendy A Bickmore

## Abstract

Long noncoding RNAs (lncRNAs) have been implicated in various biological functions including regulation of gene expression, X-inactivation, imprinting, cell proliferation and differentiation. However, the functionality of lncRNAs is not clearly understood and conflicting conclusions have often been reached when comparing different methods to investigate them. Moreover, little is known about the upstream regulation of lncRNAs. Here we show that a transcriptional co activator – PC4 and SF2 interacting protein (Psip1)/p52, which is involved in linking transcription to RNA processing, regulates the expression of the lncRNA *Hottip*. Using complementary approaches – knockdown, Cas9 mediated lncRNA deletion, analysis of lncRNA binding by Chromatin isolation by RNA purification (ChIRP) - we demonstrate that Hottip binds to the 5’ Hoxa genes located in cis, which leads to their upregulation. Moreover, the synthetic activation of *Hottip* is sufficient to induce the expression of polycomb repressed *Hox* genes in mouse embryonic stem cells (mESCs).

## Introduction

The mammalian genome encodes ~10,000 long noncoding RNAs (lncRNAs)^1^. Although very few of these molecules have been functionally characterised, a small number have been shown to function by binding to various protein complexes to regulate gene expression^2–7^. Some lncRNAs have been reported to affect gene expression *in trans^8,9^*, whereas others, such as Kcnq1ot1, Xact, Xist and Tsix, function *in cis* (reviewed in^10^). Other lncRNAs likely function in the cytoplasm through binding to other regulatory RNAs, e.g. miRNA^11^.

It has also been difficult to distinguish whether lncRNA function is conferred by the process of transcription or by the RNA molecule itself. Concerns have been raised with respect to limitations and discrepancies in various methodologies used to study lncRNA function^12^ and contrasting conclusions have often been reached when comparing knockdown and knockout studies^13–15^.

With the exception of relatively well characterized lncRNAs like Xist^16^, H19^17,18^ and Kcnq1ot1^19,20^, many recently described lncRNAs lack genetic evidence to support their function in vivo. Indeed, recent efforts to phenotype mouse knockouts for 18 lncRNA genes identified only 5 with strong phenotypes^21^. With the list of lncRNAs with unknown function increasing, there is a pressing need to rigorously dissect the functional mechanisms of individual lncRNA locus. Additionally, most research has focused on the downstream functions of lncRNAs and, with the exception of lncRNAs involved in imprinting and dosage compensation, little is known about the transcriptional regulation of lncRNAs themselves. The promoters of lncRNA genes are conserved, and are enriched for homeobox domain containing transcription factor binding sites^22^.

Mammalian *Hox* loci are important model systems for the investigation of lncRNA functions. Many noncoding RNAs within *Hox* clusters are expressed tissue specifically^23–27^, and have indeed been linked to the regulation of *Hox* mRNA genes ^8,13,28,29^ At the *Hoxa* cluster, the *lncHoxa1/Halr* - also known as *Haunt* ~ 50 kb away from 3’ end of *Hoxa* (Fig 1a) has been shown to repress *Hoxa1* expression *in cis*^30^. Importantly, a recent study demonstrated that *Haunt* lncRNA plays distinct role as a repressor while its DNA sequence functions as an enhancer for *Hoxa* genes^14^. *HOTAIRM*, located between *Hoxa1* and *Hoxa2*, is expressed antisense to coding *Hoxa* genes, and is implicated in retinoic acid induced activation of *HOXA1* and *HOXA4* during myeloid differentiation^31^. *HOTTIP* lncRNA is transcribed in an antisense direction from the 5’ end of *Hoxa13* (Fig 1a), and is reported to be important for targeting MLL through interaction with WDR5 to maintain 5’ *HOXA* expression in distal tissues^3^.

PC4 and SF2 interacting protein (Psip1) also known as LEDGF/p75 has been shown to be important for regulation of *Hox* genes in vivo^32^. We demonstrated the role of the p75 isoform of Psip1/p75 in recruiting Mll to expressed *Hox* genes^33^. The alternatively spliced short isoform of Psip1 (Psip1/p52) lacks the C-terminal Mll or integrase binding domain (IBD), but shares the chromatin binding PWWP and AT hook like domains at the N-terminus. Psip1/p52 binds to H3K36 trimethylated (H3K36me3) nucleosomes via the PWWP domain and can modulate alternative splicing by recruiting splicing factors to H3K36me3^34^.

Here, we demonstrate the role of Psip1/p52 in regulating expression of 5’ *Hoxa* genes and we show that it functions by activating expression of the Hottip lncRNA. Knockdown of either Psip1/p52 or Hottip leads to reduced expression of 5’ Hox genes - *Hoxa13, Hoxa11* and *Hoxa10*. Using CRISPR-Cas9 mediated deletion versus dCas9-VP160 mediated transcriptional activation, we demonstrate the function of Hottip lncRNA transcription in *Hox* regulation and the direct role of Psip1/p52 in activating lncRNA Hottip, which further maintains 5’ *Hoxa* expression.

## Results

### Psip1 is required for expression of lncRNA Hottip

In mammals, active *Hox* genes are maintained by complexes containing the MLL (Mix lineage leukemia) histone H3K4 methyltransferases, and repression is maintained by Polycomb (PcG) complexes^35^. We recently demonstrated a role for Psip1/p75 in recruiting Mll to expressed genes from the 5’ end of Hoxa^33^. Most strikingly, at the extreme 5’ end of *Hoxa*, beyond *Hoxa13*, where the Hottip lncRNA^3^ is located, Mll1 binding is completely lost in *Psip1^−/−^* MEFs compared to wild type (WT), this is accompanied by a concurrent loss of H3K4me3 and Menin - a common component of Mll1 and Mll2^36^ (Fig. 1a). Loss of Psip1 also results in a complete loss of *Hottip* expression and reduced expression of *Hoxa13*, which has been described as a target of Hottip (Fig. 1b)^3^. In contrast, other previously described Hottip target genes - *Hoxa9, a10*, and *a11*^3^ are up-regulated in *Psip1^−/−^*MEFs despite the loss of *Hottip* expression ^33^. Nascent run-on analysis shows that these effects occur at the level of transcription (Fig. 1c). Together with the binding of Psip1 to the 5’ *Hoxa* cluster (Fig 1a), this suggests that Psip1 has a direct role in regulating expression of *Hottip* lncRNA. Psip1 is also highly expressed in distal limb buds of mouse embryos, where *Hottip* and 5’ *Hox* genes are expressed (Fig. S1a)^3^.

**Figure 1.**
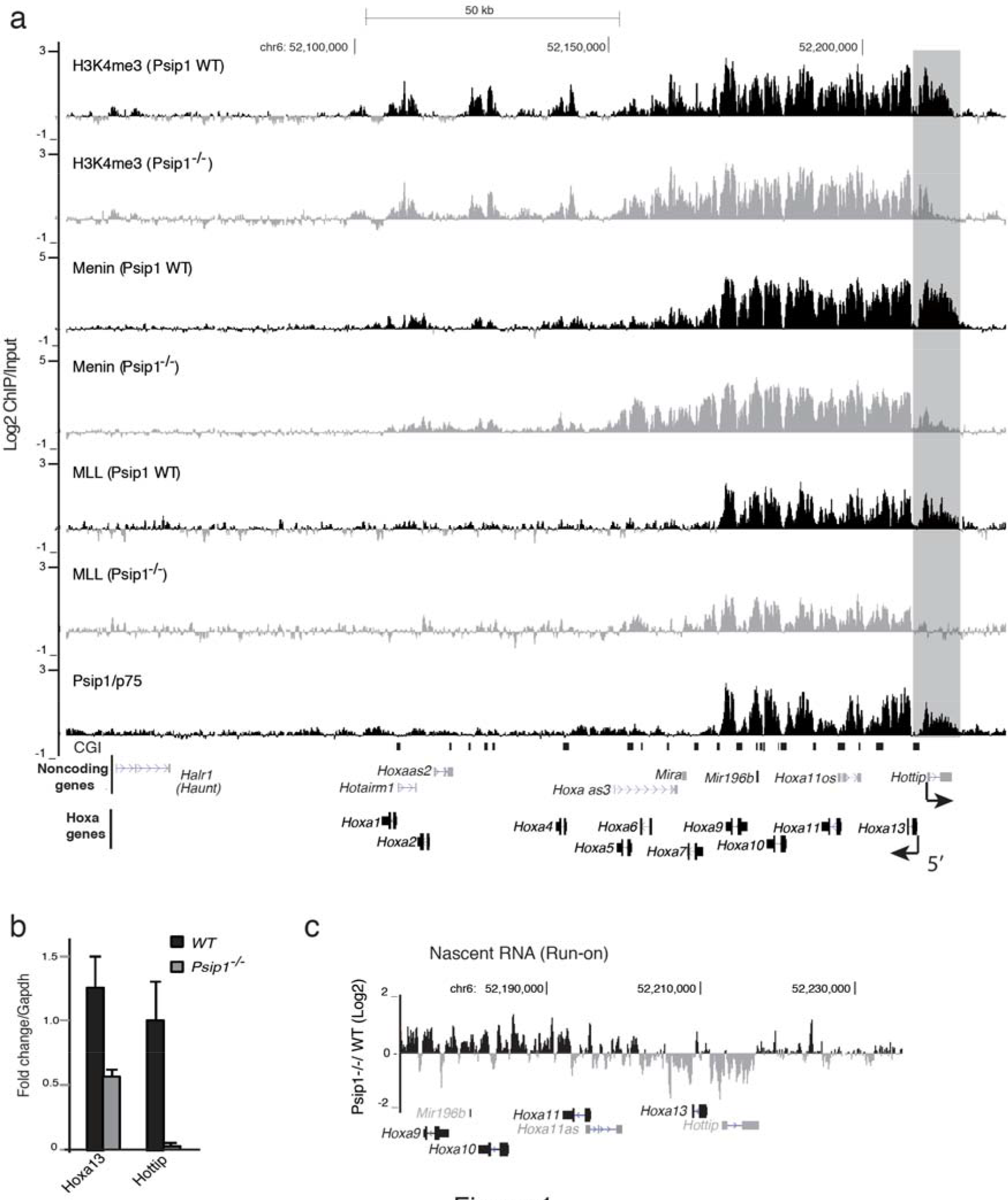
Reduced Hottip expression and Mll occupancy in *Psip1−/−*. a) Mean Log2 ChIP/input for Psip1/p75, Mll1, Menin and H3K4me3 in WT and *Psip1−/−* MEFs over *Hoxa* clusters from custom tiling arrays^33^. Annotated noncoding transcripts (grey, top) and *Hox* gene transcripts (black) are shown below. (n = 2 biological replicates). Genome co-ordinates are from the mm9 assembly of the mouse genome. Direction of transcription for *Hoxa13* and *Hottip* genes are indicated below. b) Mean (± s.e.m) expression, assayed by RT-qPCR and normalized to Gapdh, of *Hoxa13* and *Hottip* in WT and *Psip1−/−* MEFs, (n = 3 biological replicates). c) Log2 ratio of *Psip1−/− / WT* run-on transcribed RNA (nascent RNA) over 5’ *Hoxa* genes.

### Psip1/p52 and Hottip are essential for the maintenance of 5’ *Hoxa* gene expression

To identify which isoform of Psip1 regulates Hottip, two independent lentiviral shRNAs specifically targeting the 3’ UTR of p52, C-terminus of p75 and Hottip RNA
were transduced into a limb mesenchymal cell line (14FP) which retains the distal limb-specific expression pattern of *Hox* genes^37^. Knockdown efficiency was confirmed by RT-qPCR analysis (Fig. 2a) and by immunoblotting for Psip1 isoforms (Fig. 2b). Upon specific knockdown of Psip1/p52, expression of 5’ *Hoxa* genes - *Hoxa13, all* and *a10* is down regulated with the strongest abrogation of *Hoxa13* mRNA (Fig. 2a). Knockdown of p52 had a strikingly similar effect on 5’ *Hoxa* expression as knockdown of *Hottip* (Fig. 2a), a result consistent with the knockdown of human HOTTIP in human foreskin fibroblasts^3^. Knockdown of Psip1/p75 does not affect 5’ *Hoxa* genes in these cells, which demonstrates that it is the p52 isoform of Psip1, not p75 that specifically activates *Hottip* lncRNA transcription. Moreover, these data indicates that the Hottip lncRNA is involved in maintaining the active chromatin domain at 5’ *Hox* genes^3^. Knockdown of *p52, p75* and *Hottip* using independent shRNAs confirms that mis-regulation of *Hox* genes is not due to off-target effect of shRNAs (Supplementary Figure 1b).

We found a significant reduction in Hottip nascent RNA levels in the p52 knockdown cells (Fig. 2a), which shows that reduced *Hottip* RNA levels are not simply due to the known effect of Psip1/p52 on RNA splicing^34^. Moreover, reduced Hottip expression in p52 knockdown cells (shRNA targeting 3’ UTR of p52) could be rescued by transducing an shRNA resistant p52 cDNA (Fig. 2c).

**Figure 2.**
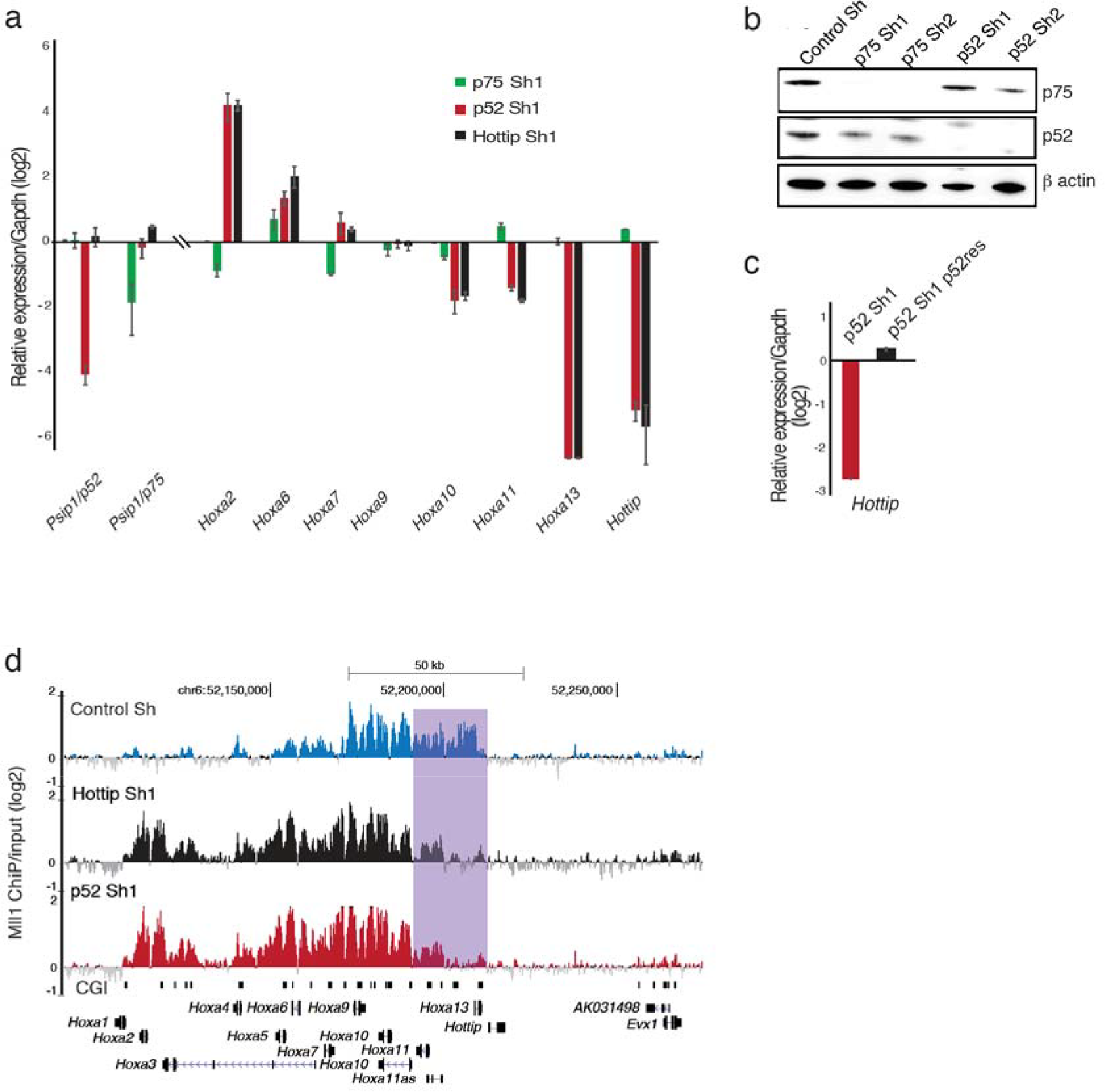
Psip1/p52 and Hottip are important for expression of 5’ Hoxa genes. a) Log2 mean (± s.e.m) expression, assayed by RT-qPCR and normalized to Gapdh, of Hoxa genes, along with *Psip1/p52*, *Psip1/p75*, and *Hottip* RNA, in limb cells transduced with shRNAs targeting p52 (red bars, p52 sh1) p75 (green bars, p75 sh1) and Hottip (black bars, Hottip sh1) relative to cells transduced with a mammalian non-targeting sh RNA (control) (n = 3 biological replicates). b) Immunoblotting of limb cells after shRNA knockdown of p52 and p75 Psip1 isoforms with Psip1 antibody (A300-847a) which recognizes both p52 and p75^34^. β-actin served as loading control. Two different sets of shRNAs (sh1 in (a) and sh2 in Supplementary Fig. 1) were used for knockdown along with a mammalian nontargeting shRNA as control (control_sh). c) Mean log2 expression of *Hottip*, in limb cells transduced with sh RNAs targeting p52 (red bars, p52 sh1) and those cells rescued transiently with shRNA resistant p52 cDNA (green bars, p52 sh1 p52res). Fold change in expression was normalized to Gapdh, relative to mammalian non-targeting shRNA (control) (n = 3 biological replicates). d) Mean Log2 ChIP/input across *Hoxa* cluster for Mll1 from limb cells transduced with control shRNA (Control_Sh), shRNA targeting p52 (p52 sh1) and Hottip (Hottip sh1) as described in (a) & (b). Genome co-ordinates are from the mm9 assembly of the mouse genome.

### Mll1 occupancy over *Hoxa* cluster is altered upon *p52* & *Hottip* knockdown

It has been suggested that Hottip has a role in maintaining an MLL complex through interaction with the WDR5 component^3^. Consistent with this, Mll1 occupancy was significantly reduced across 5’ *Hoxa* genes upon knockdown of *p52* or *Hottip* compared to control knockdown (Fig. 2d), Intriguingly, upon depletion of p52 (Fig. 2a and b) and Hottip (Fig. 2a), Mll1 was redistributed to 3’ *Hoxa* genes, which is consistent with the increase in expression of 3’ *Hoxa* genes upon p52 or *Hottip* depletion (Fig. 2d). Loss of Mll1 upon p52 and Hottip knockdown is consistent with the reduced Hottip expression along with reduced Mll1, H3K4me3 and Menin across *Hoxa13* and *Hottip* loci in *Psip1^−/−^*MEFs (Fig. 1a).

### Deletion of *Hottip* leads to reduced expression of 5’ *Hoxa genes*

Most lncRNA depletion studies are done by si/sh RNA mediated knockdown, but the conclusions reached have often been different from those after genetic deletion of the loci encoding the lncRNAs^13–15^. We therefore used two pairs of guide RNAs with cas9 nickase (cas9n) to delete the gene body of *Hottip* (HottipΔ) in limb mesenchymal cells, leaving the promoter intact (Fig. 3a). RT-qPCR of *Hoxa* genes in homozygous HottipΔ cells showed a >50% reduction in expression of *Hoxa13* and *all* (Fig. 3b). Similar to Psip1 and *Hottip* knock down studies (Fig. 2a), expression of 3’ *Hoxa* genes, such *Hoxa2, a6* and *a7* increased compared to WT cells.

### Hottip RNA is localized at 5’ *Hoxa* genes

To find the direct targets of Hottip lncRNA in limb cells, chromatin isolation by RNA purification (ChIRP)^38^ was performed with multiple antisense oligo pools for Hottip. RT-qPCR analysis of ChIRPed RNA showed specific enrichment for *Hottip* (Fig. 3c). qPCR analysis of Hottip ChIRPed DNA shows specific enrichment of *Hottip* RNA over the promoters of *Hoxa13, all* and *a10* (Fig. 3d). This was abolished in *HottipΔ* cells demonstrating the specificity of the *Hottip* ChIRP (Fig. 3b). *Hottip* was undetectable across 3’ *Hoxa* genes (*a9, a7, a1*), which are nonetheless upregulated in *HottipΔ* cells (Fig. 2), demonstrating that misregulation of 3’ *Hoxa* genes is a secondary event which does not involve direct binding of *Hottip*.

### Induction of Hottip lncRNA is sufficient to activate 5’ *Hoxa* genes

It was possible that reduced expression of 5’ *Hox* genes in *HottipΔ* cells is due to loss of *cis* regulatory elements located within the deleted region, rather than due to loss of the *Hottip* RNA, to rule out this possibility we synthetically activated Hottip expression in mES cells, by recruiting a catalytically inactive Cas9 (dcas9) fused to a synthetic activator - VP64/VP160 dcas9-VP160 (VP16 x10)^39–42^. CRISPR dCas9 mediated transcriptional activation has been shown to be ineffective when guides are targeted >1kb from TSS^39^. Unlike human HOTTIP which is transcribed bidirectionally from the *HOXA13* CpG island promoter (Fig. 4a)^3^, the Mouse *Hottip* promoter is ~2 kb away from the TSS of *Hoxa13* which allowed us to recruit dcas9-VP160 to the promoter of *Hottip* and not to that of *Hoxa13* (Fig. 4a).

**Figure 3.**
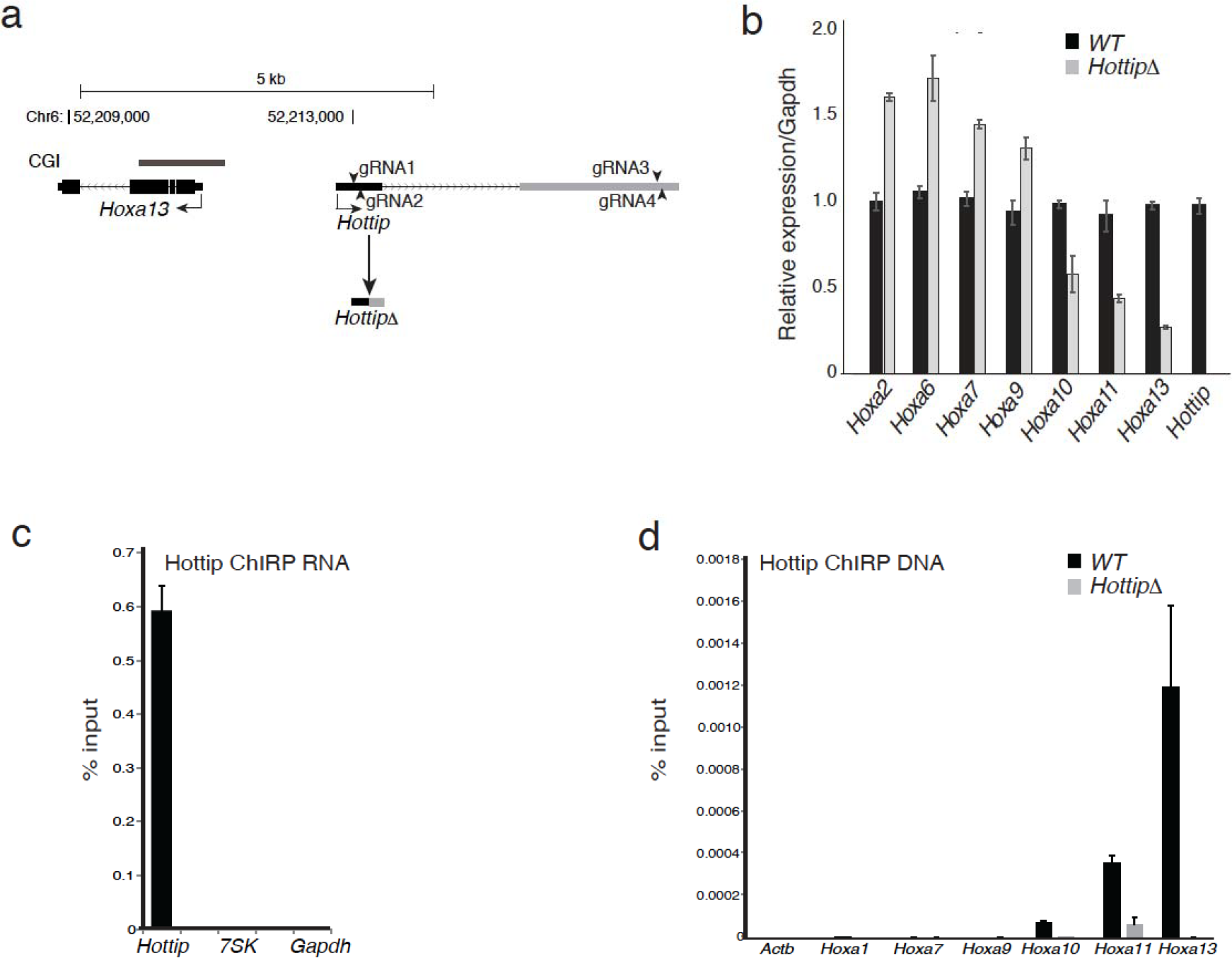
a) Schematics showing the mouse *Hoxa13* and *Hottip* loci. The CpG Island (CGI) at the *Hoxa13* promoter is shown in a gray bar. Genome co-ordinates are from the mm9 assembly of the mouse genome. Guide RNA binding sites for deletion of *Hottip* are shown as arrow heads. The deletion product of *Hottip* (*HottipΔ*) is shown below. b) Mean (± s.e.m) expression, assayed by RT-qPCR and normalized to *Gapdh*, of *Hoxa* genes and *Hottip*, in wild-type (black bars, WT), and *Hottip* knock out (gray bars, *HottipΔ*) limb mesenchymal cells, (n = 3 biological replicates). c) RT-qPCR showing mean (± s.e.m) ± percentage (%) enrichment over input for *Hottip*, *7SK* and *Gapdh* RNAs from Hottip ChIRP pulldown from two experiments. d) qPCR showing mean (± s.e.m) percentage (%) enrichment over input of ChIRPed DNA at promoters of *Actb*, *Hoxa1*, *Hoxa7*, *Hoxa9*, *Hoxa10*, *Hoxa11*, and *Hoxa13* from Hottip ChIRP experiments in wild type (black bars, WT) and Hottip knock out limb mesenchymal cells (grey bars, *HottipΔ*).

Microarray and RT-qPCR analysis showed specific upregulation of 5’ (*a13, a11* and *a10*), but not 3’, *Hoxa* genes upon dcas9-VP160 mediated Hottip activation (Fig. 4b & 4c). In contrast, specific recruitment of dCas9-VP160 to the *Hoxa13* promoter led to up regulation of only *Hoxa13* - expression of other *Hoxa* genes was unaltered (Fig. 4b). Recruitment of dCas9-VP160 to the *Hottip* promoter in *HottipΔ* limb cells led to only a modest upregulation of *Hoxa13* compared to WT cells (Fig. 4c), pointing to the importance of *Hottip* RNA in the regulation of *Hoxa* genes.

**Figure 4.**
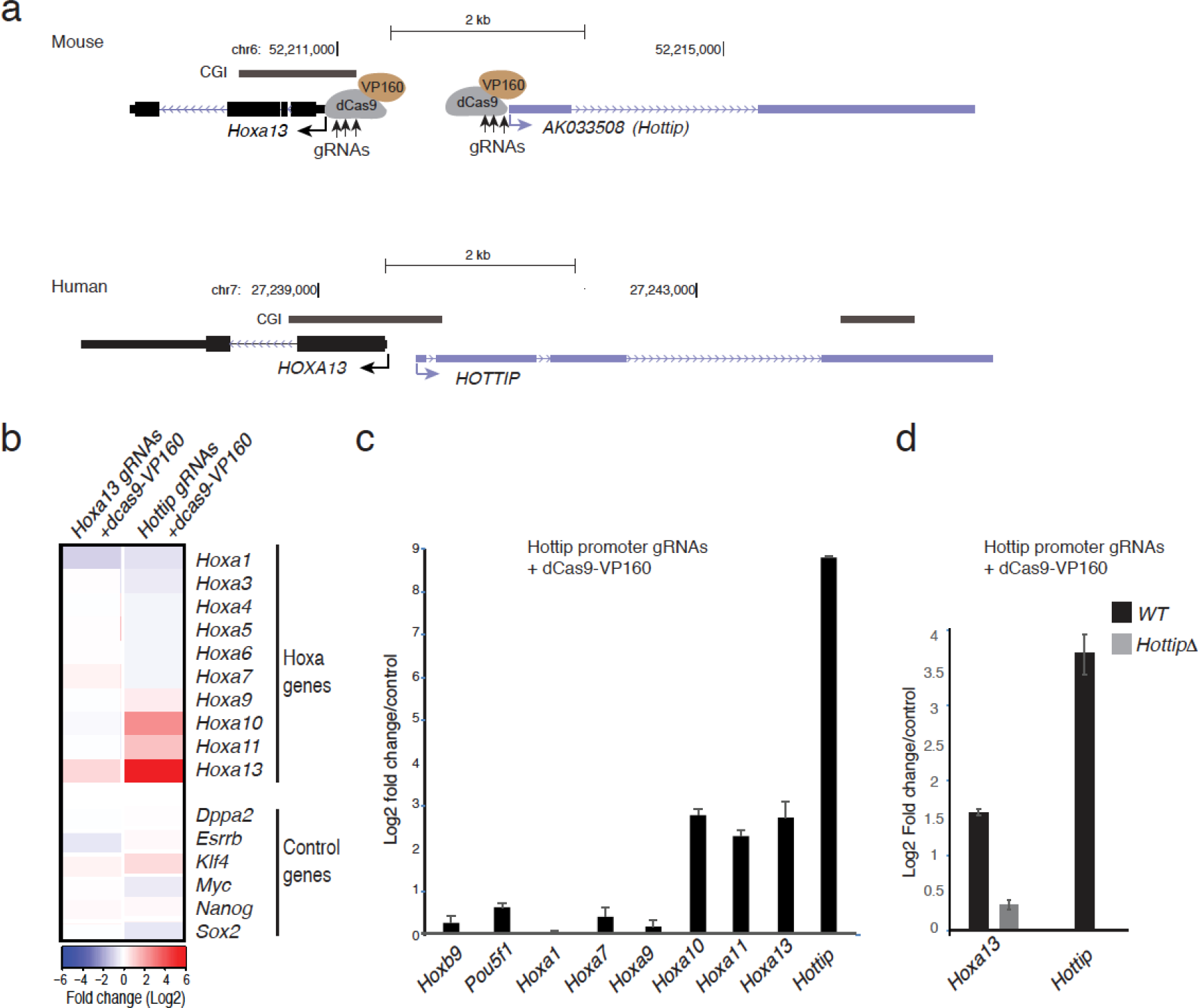
a) Schematics showing UCSC genomic coordinates of *Hottip* and *Hoxa13* and CpG Islands (CGI) in the mouse (top, mm9) and human (bottom, hg19) genomes. Schematics of guide RNA mediated recruitment of dCas9-VP160 to the *Hottip* promoter is also shown. Direction of transcription is indicated. b) RT-qPCR data showing mean (± s.e.m) log2 fold change in expression of *Hottip*, *Hoxa13*, *a11*, *a10*, *a9*, *a7*, *a1*, *Pou5f1* and *Hoxb9* upon guide RNA mediated recruitment of dCas9-VP160 to the *Hottip* promoter (a) in mouse ES cells. Data were normalized to those from a dcas9 control (n= 3 biological replicates) c) Similar to (b) heat map showing the log2 mean fold change in expression of *Hoxa* and pluripotency associated genes (control genes) from an expression microarray, upon co-transfection of guide RNAs recognizing the *Hottip* promoter (Hottip gRNAs + dcas9-VP160). dCas9-VP160 was also co transfected with guide-RNAs recognizing *Hoxa13* promoter (Hoxa13 gRNAs + dcas9-VP160). d) Similar to (b) mean log2 fold change in *Hottip* and *Hoxa13* expression in wild type ES cells co-transfected with guide-RNAs recognizing the *Hottip* promoter and dCas9-VP160 (Black bars, WT). *Hoxa13* expression in *Hottip* knock out limb mesenchymal cells is also shown (grey bar, *HottipΔ*).

## Discussion

Psip isoforms regulate the expression of *Hox* genes through two distinct mechanisms. We have previously shown that the longer p75 isoform binds directly to Mll through its MLL or integrase binding domain (IBD) and recruits Mll to active *Hox* clusters^33^. Here we have demonstrated that the shorter isoform Psip1/p52 - that lacks the C-terminal Mll1 binding domain of p75 - controls 5’ *Hoxa* genes by activating the expression of *Hottip* lncRNA.

With the recent increase in the number of lncRNAs identified using high-throughput sequencing technologies, the list of these transcripts with unknown upstream transcriptional regulation and downstream functional mechanism is growing. Till date only few proteins have been shown to transcriptionally regulate lncRNAs - Oct4 and Nanog have been shown to directly regulate expression of several lncRNAs in mESCs^43^ and p53 induces expression of several lncRNAs including lincRNA-p21^44^. Recently, two micro RNAs miR-192, miR-204 have been demonstrated to post transcriptionally silence the HOTTIP lncRNA, which leads to the reduced viability of Hepatocellular carcinoma (HCC) cells^45^.

Although transcription at noncoding enhancer elements has been associated with enhancer activity^46,47^. Exosome mediated degradation of enhancer transcripts suggests that in most of the cases the process of transcription itself could be sufficient but not functional RNAs^48,49^. Enhancer like function of lncRNAs has been demonstrated in only few cases including HOTTIP^3^.

Together, we demonstrate the specific transcriptional regulation of *Hottip* by a transcriptional coactivator Psip1/p52, which has been shown to have higher in vitro coactivator activity than Psip1/p75 isoform^50^. Increasing evidences shows the existence of specifically regulated lncRNAs at transcriptional and post-transcriptional level. Further research is needed to distinguish the importance of specifically regulated lncRNAs from enhancer like function of the DNA elements at these loci in *cis* regulation of gene expression.

## Acknowledgements

We thank Prof. Alan Engelman (Harvard Medical School) for *Psip1^−/−^* MEFs and the Psip1/p52 and Psip1/p75 retroviral rescue plasmids. We thank Prof. Robert Hill (MRC-IGMM, University of Edinburgh) for 14FP cells. This work was funded by Medical Research Council UK and Wellcome Trust (WT085767).

## Methods

### Cell lines

*Psip1^−/−^* and its corresponding WT MEFs^33,51^ were kind gift of Prof. Alan Engelman (Dana-Farber Cancer Institute, USA). Limb mesenchymal cells (14FP) isolated from the posterior mesenchyme of E11.5 mouse embryos from an Immortomouse (H-2kb-tsA58) × CD1 cross, are as previously described^37^ and were a kind gift from Robert Hill (University of Edinburgh).

### shRNA knockdown

Lentiviral shRNAs (pLKO.1 vectors) targeting Psip1/p52, Psip1/p75 and Hottip (Supplementary Table 1) were transduced as described by the manufacturer (Sigma Aldrich).

### RTqPCR

cDNA prepared from superscript II reverse transcriptase was used to perform expression qPCR. All qPCRs were performed with three biological replicates in a LightCyler 480 (LC480, Roche), and the data was normalized to *Gapdh*. Details of the oligos are given in the Supplementary Table 2

### Chromatin Isolation by RNA Purification (ChIRP)

Anti-sense oligo probes tiling the mouse Hottip RNA were designed using the web tool from Stellaris FISH Probe Designer(https://www.biosearchtech.com/support/education/stellaris-rna-fish) Biosearch Technologies, CA, USA). Biotinylated oligos were synthesized by Sigma-Aldrich. ChIRP was performed in limb mesenchymal cells as described previously^38^. RNA was isolated from 20% of the ChIRPed beads and used for RT-qPCR for Hottip, 7sk and Gapdh specific primers and rest was used to perform qPCR for Hoxa genes. Primer details are given in Supplementary Table 2.

### RNA In situ

RNA in situ hybridization for *mHottip* in 10.5 days mouse embryos were performed as previously described^52^. Details of oligos used to PCR amplify Hottip cDNA including T7 (sense) and T3 (Antisense) promoter sequences are given in Supplementary Table 3.

### ChIP, antibodies and data analysis

ChIP was performed as described previously^33^, using antibodies for Mll1 (Active Motif 61295, 61296), ChIP DNA was hybridized to a custom *Hox* array and data was normalized as described previously^33^.

### CRISPR mediated deletion of Hottip

Guide RNAs were designed using Zhang laboratory web tool (http://crispr.mit.edu). Paired guide RNAs (gRNAs) (Supplementary Table 4) were designed to target the murine *Hottip* genomic locus beyond the transcription start site (TSS) and before the transcription end site (Figure 2e). gRNAs were cloned into the D10A nickase mutant version of cas9 (cas9n) containing pSpCas9n(BB)-2A-GFP (PX461)^53^. A pool of four gRNA containing plasmids were transfected into limb cells using FuGENE HD transfection reagent and FACS sorted 48 hours after transfection for GFP+ cells. Homozygous deletion of Hottip was confirmed by PCR and Sanger sequencing, primers used are given in Supplementary Table 2.

### Cas9-mediated activation of *Hottip* and *Hoxa13*

Guide RNAs targeting the promoters of *Hottip* and *Hoxa13* (Supplementary Table 5) were designed as above and cloned into pSLQ137 1 ^49,54^. gRNA containing plasmids encoding mCherry and puromycin resistance were co-transfected with dCas9-VP160^42^. 24hrs after transfection, transfected cells were selected by addition of 2μg/ml puromycin for another 24 hrs. RNA was extracted 48hrs after transfection for RT-qPCR and microarray gene expression analysis was performed according to the manufacturer’s protocol (Agilent Technologies). To investigate whether activation of lncRNA Hottip is sufficient to induce the expression of target 5’ *Hoxa* genes, 5 guide RNAs targeting the *Hottip* and *Hoxa13* promoters were cotransfected with dCas9-VP160 in mESCs (Supplementary Table 5). Cells were harvested 48hrs after transfection for both RNA expression analyses by RT-qPCR and expression microarrays. Plasmids containing non-targeting guide RNAs and dCas9 alone (dcas9Δ) served as controls.

